# Unlocking data in *Klebsiella* lysogens to predict capsular type-specificity of phage depolymerases

**DOI:** 10.1101/2024.07.24.604748

**Authors:** Robby Concha-Eloko, Beatriz Beamud, Pilar Domingo-Calap, Rafael Sanjuán

## Abstract

Viral entry is a critical step in the infection process. *Klebsiella* spp. and other clinically relevant bacteria often express a complex polysaccharide capsule that acts as a barrier to phage entry. In turn, most *Klebsiella* phages encode depolymerases for capsule removal. This virus-host arms race has led to extensive genetic diversity in both capsules and depolymerases, complicating our ability to understand their interaction. This study exploits the information encoded in *Klebsiella* prophages to model the interplay between the bacteria, the prophages, and their depolymerases, using a graph neural network and a sequence clustering-based method. Both approaches showed significant predictive ability for prophages capsular tropism and, importantly, were transferrable to lytic phages. In addition to creating a comprehensive database linking depolymerase sequences to their specific targets, this study demonstrates the predictability of phage-host interactions at the subspecies level, providing new insights for improving the therapeutic and industrial applicability of phages.

## Introduction

In bacteriophages and other viruses, recognition of the host is considered a critical step for successful infection (Silva, Storms, and Sauvageau 2016). This process is mediated by virus-encoded receptor binding proteins (RBPs) and mostly comprises two categories of phage proteins: tail fiber and tail spike proteins (or depolymerases). Tail fiber proteins are generally non-enzymatic and bind to a protein receptor on host surface (Taslem Mourosi et al. 2022). In contrast, tail spikes are characterized by the presence of a polysaccharide-degrading domain, sometimes referred to as depolymerase domain (Pires et al. 2016; Latka et al. 2019). The depolymerase domain can adopt a variety of fold conformations (Lombard et al. 2010; Drula et al. 2022; Concha-Eloko et al. 2024) and is responsible for the specific recognition and degradation of complex sugars like exopolysaccharides (EPS), capsular polysaccharides (CPS) or lipopolysaccharide (LPS) (Topka-Bielecka et al. 2021). Current models describe phage-bacteria interactions mediated by the depolymerase domain in two steps: (i) reversible binding to the capsule as the primary receptor through the depolymerase domain before its degradation, and (ii) subsequent access to the secondary receptor located on the membrane surface before entry into the host (Dunstan et al. 2021).

Phages are particularly relevant against ESKAPE pathogens, which are regarded as being among the most urgent threats to global health (Naghavi et al. 2024). This group includes *Klebsiella pneumoniae*, a gram-negative bacterium that expresses a capsule that provides protection against external stressors such as the immune system, antibiotics, and phages (Clegg and Murphy 2016). The capsule of *Klebsiella* spp., encoded by the Wzy system, is highly diverse (Yun Yang et al. 2021). Seventy-seven serological types have been described, and more recently, 180 K-loci (or KL types) have been identified based on analysis of the K-locus, the coding region involved in capsule synthesis (Lam et al. 2022, 2). The KL type is the most determining factor of infectivity for most phages targeting *Klebsiella* (de Sousa et al. 2020; Beamud et al. 2023; Ferriol-González et al. 2024), but our ability to determine the KL tropism of a given phage based on depolymerase sequence alone remains limited. Through depolymerase activity, phages apply selective pressure on bacteria, sparking an evolutionary arms race. On one hand, bacteria can evade phage infection through mutations encoding morphological changes in their capsule or the complete loss of capsule expression (Fang et al. 2022; Rendueles, De Sousa, and Rocha 2023; Tang et al. 2023). Capsule loss and reacquisition is often driven by horizontal gene transfer, which can result in rapid KL type shifts (Haudiquet et al. 2021). On the other hand, phages counter bacterial evasion through depolymerase domain evolution, primarily via horizontal gene transfer across diverse phages (Pas et al. 2023; Yiyan Yang et al. 2024), leading to a poor correlation between genome-based phage phylogenies and KL tropism (Beamud et al. 2023). These processes contribute to the significant diversity observed in depolymerase sequences (Oliveira, Drulis-Kawa, and Azeredo 2022).

Recent advances in the field of machine learning have paved the way for addressing important questions related to phages. Many of these advancements have been grounded in new paradigms, such as the attention mechanism, which can be defined as the ability of a model to focus on specific parts of the input to enhance its predictive performance (Bahdanau, Cho, and Bengio 2016; Vaswani et al. 2023). This principle allowed the emergence of the transformer architecture, whose application in biology led to the development of Protein Language Models (PLM) (Devlin et al. 2019). PMLs, notably Evolutionary Scale Modeling 2 (ESM2) recently developed by MetaAI, were trained on millions of protein sequences. The objective of training was to predict the identity of a token (*i.e* one or few consecutive amino acids) randomly concealed within a given protein sequence (Ferruz, Schmidt, and Höcker 2022). Through this self-supervised approach, PLMs can capture complex patterns and relationships between amino acids in protein sequence into embedded representations. These embeddings consist of feature vectors that encapsulate biochemical and biophysical properties of the protein sequence (Lin et al. 2023). Various studies have reported models that use embedding representations for downstream tasks such as sequence or token classification, with remarkable results (Kang et al. 2022; Sledzieski et al. 2023). Over the past few years, several teams have attempted to model phage-bacteria interactions and predict their outcomes. Models have been successfully generated to make predictions at the genus and species levels (Boeckaerts et al. 2021; Shang and Sun 2022; J. Pan et al. 2023; Roux et al. 2023). However, in some specific settings, such as the therapeutic one, these levels of resolution are not satisfactory. Indeed, phages are highly specific and typically infect a few strains within a species (P. Domingo-Calap, Georgel, and Bahram 2016; de Jonge et al. 2019). Three aspects are crucial for attaining a higher level of resolution when predicting the outcome of phage-bacteria interactions: (i) extensive high-quality training data, (ii) strong biological premises and (iii) state-of-the-art machine learning techniques. Recent studies (Gaborieau et al. 2023; Briers et al. 2023) managed to make great progress in this direction, but still reported limitations related to the a lack of training data.

In this work, the challenge of the amount of training data was addressed by leveraging the information contained in *Klebsiella* lysogens and their prophages. Large numbers of depolymerase domain sequences encoded by prophages were retrieved with the aim to generate a model predicting the outcome of the interaction between a KL type and a phage given its depolymerase domain sequences. Two approaches were used in this study. Firstly, a graph-based approach was developed to model the complex interplay between the depolymerase domain, prophages, and bacteria. Embedding representations were generated for each depolymerase domain sequence and aggregated using a graph convolution network relying on the attention mechanism (GATv2) (Brody, Alon, and Yahav 2022). The resulting representation was then fed into a deep neural network for binary classification. Secondly, a sequence-clustering-based model employing categorical features, corresponding to the presence-absence of representative depolymerase domains, was used to fit the Random Forest algorithm. Both approaches displayed significant predictive abilities for prophages, which could be transferred to lytic phages. Moreover, these approaches have allowed the generation of a database of depolymerase domain sequences linked to their target KL types. This contributes to the understanding of phage-bacteria interactions mediated by the depolymerase domain sequence and capsule and offers potential applications in therapeutic and industrial settings.

## Material and Methods

### Bacterial genomes and capsule typing

The genomes of *Klebsiella* strains were downloaded from the NCBI Reference Sequence *Database* through the PanACoTa package (Perrin and Rocha 2020). In order to retrieve all the strains from the Genus *Klebsiella*, the “prepare” command with the “-T 570” parameter were used. The K-loci type (KL type) of the bacterial strains were then predicted using Kleborate (https://github.com/klebgenomics/kleborate) (Lam et al. 2022). The strains that presented a level of confidence of “perfect”, “very high”, “high” and “good” were kept for the analysis. Finally, the genomes were annotated with Prokka v.1.13 (Seemann 2014).

### Phylogenetic tree

The PanACoTa pipeline was used in a similar manner as used by Haudiquet et al. 2021. Briefly, the set of genes present in the genomes of the dataset (or pan-genome) was determined with the “pangenome” command. The output was then used to compute the set of genes present in more than 99% of the genomes in the dataset (or core-genome) using the “corepers” command with the “-t 0.99” parameter. Subsequently, the genes from the core-genome were aligned using the “align” command. The aligned nucleotide sequences of each genes were then concatenated, resulting in the alignment of nucleotide sequences of 498,179 bp. Phylogenetic inferences were executed with IQ-TREE (v.2.1.4-beta COVID-edition) using the *ModelFinder Plus* parameter “-m MFP”, with 1000 replicates for ultrafast bootstrap “-B 1000” (Minh et al. 2020). The resulting best model was the (GTR+F+I) corresponding to the general time-reversible model with empirical base frequencies allowing a proportion of invariable sites. The tree was well supported with an average ultrafast bootstrap value of 99.85%.

### Prophage prediction

Prophages were predicted using the PhageBoost program based on an machine learning approach (Sirén et al. 2021) trained to distinguish the viral signal from the background signal of a bacterial genome, based on 1,587 different features. A threshold of 0.70 confidence was applied, allowing the number of called prophages to be in concordance with the reported mean number of prophages in *Klebsiella* spp. in previous studies (de Sousa et al. 2020). In order to identify prophages belonging to the same strain, the pairwise average nucleotide identity (ANI) was computed between the prophages from the dataset using FastANI (Jain et al. 2018). The ANI is defined as the mean nucleotide identity of orthologous gene pairs shared between two genomes. Two prophages were considered from the same strain if the calculated ANI>99% with a bi-directional coverage >80%.

### Recovering the KL type of the infected ancestral bacteria

To limit the bias induced by the horizontal gene transfers of the K-locus, the KL type of the bacteria before phage infection, *i.e* of the infected ancestor, was investigated for each prophage. First, the states of the ancestors of the bacteria from the dataset were inferred using the phylogenetic tree and the KL types computed for each strain with PastML (v1.9.34) (Ishikawa et al. 2019). The maximum likelihood algorithm MPPA with the recommended character evolution model F81 was used, allowing the attribution of multiple states when they had similar and high probabilities. Next, the infected ancestor was identified for each prophage through the Bio.Phylo package (Talevich et al. 2012) using a tailored algorithm (**Supplementary Algorithm 1**). The whole process is summarized in **Figure 1A**.

**Figure 1.**
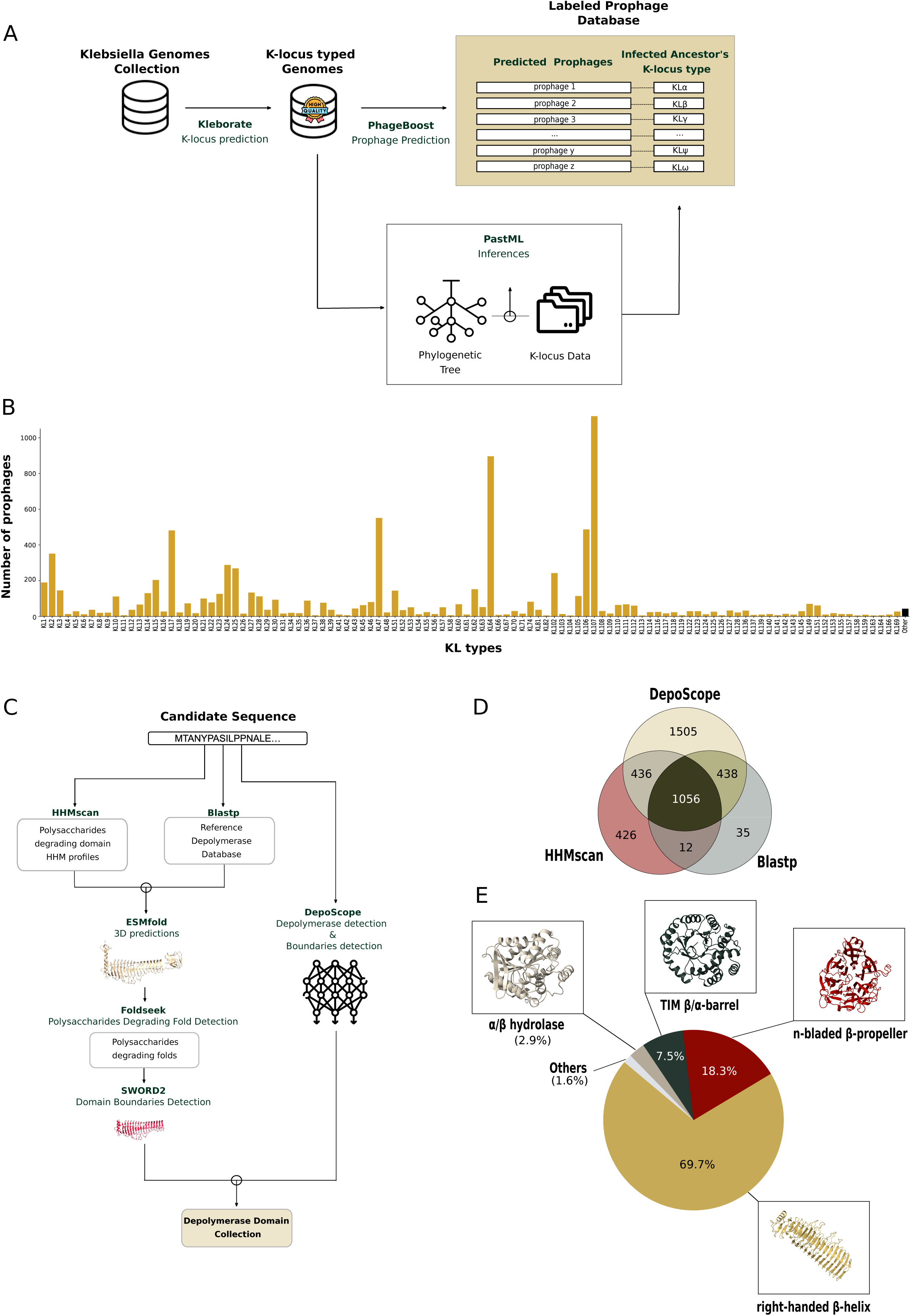
Generation of prophage collection with the KL types of the infected ancestors, along with their encoded depolymerase domains **A:** Schematic representation of the pipeline employed to generate the collection of labeled prophages from the *Klebsiella* genome dataset. **B**: Bar plot of the number of prophages carrying at latest one depolymerase on the y-axis across each KL types represented on the x-axis. The KL types represented by less than 5 prophages were assigned to the “Other” category. **C:** Pipeline for the identification of depolymerase domain sequences from prophage proteomes. **D:** Venn diagram of the depolymerase domain identified through the 3 methods employed. They involved the deep learning based method Deposcope, a method based on HMM profiles of proteins with depolymerase polysaccharides degrading activity using HHMscan and blastp to scan the proteins against a collection of characterized depolymerases from the literature. **E:** Proportions of the depolymerase folds identified.

### Identification of the depolymerase domain sequences

Phage sequences were annotated with Prodigal from Pyrodigal v.0.2.1 (https://github.com/althonos/pyrodigal), integrated in Phageboost. Sequences with a length inferior to 200 amino acids were discarded. Three different approaches were employed to identify the depolymerase domain sequences in the prophages. The first approach leveraged a database of Hidden Markov Models (HMM) profiles of domains associated with a polysaccharides degrading (PD) activity (Concha-Eloko et al. 2024). Multiple sequence alignments (MSA) were built around the candidate protein sequences using MMseqs (Steinegger and Söding 2017) along the UniRef90 database. The resulting MSAs were then scanned against the HMM profiles of the PD domains using the HHsearch (Steinegger et al. 2019). The hits with a score exceeding 20 and an alignment spanning at least 30 amino acids were considered candidate depolymerases. In the second approach, prophage protein sequences were directly screened against a database of 333 depolymerase proteins from previously described phages (Beamud et al. 2023) using blastp (Altschul et al. 1990). The hits presenting a bitscore exceeding 75 were added to the set of candidate depolymerase. Subsequently, the candidate depolymerase set resulting from the two approaches above were processed jointly. The 3D structures of the candidate depolymerases were predicted using ESMfold (Lin et al. 2023). Then, the predicted structures were scanned using FoldSeek (Van Kempen et al. 2023) against a database of 3D domain folds involved in a PD activity. The folds represented in the database included the α/α toroid, the right-handed β-helix, the TIM β/α-barrel, the n-bladed β-propeller (with at least 4 antiparallel β-strands), the flavodoxin-like fold and the α/β hydrolase fold. The hits that presented a probability of being a true positive greater than 0.5, or 0.2 when the query was the right-handed β-helix, were considered depolymerase proteins. Next, the 3D structure of the depolymerase proteins were dissected into protein units using SWORD2 (Cretin et al. 2022). These protein units were scanned against the database of 3D structures of PD domains to infer the definitive depolymerase domain from the best match. The third approach was based on the deep learning model DepoScope (Concha-Eloko et al. 2024). This model performs a binary classification of proteins for the identification of depolymerase proteins, as well as the delineation of the catalytic domain. The final depolymerase domain database was generated by sequences that were identified by at least one of the methods. An overview of the subsequent data processing is given in **Figure 1C**.

### Graph-based approach for modeling prophage-CPS Interactions

Graphs can represent complex relationships between objects using nodes to encode the objects and edges to encode their connections (Veličković 2023). Here, a heterogeneous graph was designed, with three node types: the infected bacteria (nodes A), the prophages (nodes B1) and the depolymerases (nodes B2). These nodes were connected with two types of directed edges: the edges (B2→B1) connecting the depolymerase to the prophage where it was identified, and the edges (B1→A) connecting the prophage to the corresponding infected bacteria. To each node type were associated specific features: the nodes A were characterized by the KL type of the infected ancestor; the nodes B1 had no initial specific feature; and the nodes B2 were characterized by the 1,280 dimensions embedding representations of the depolymerase domain’s amino acid sequence, computed using the ESM2 model esm2_t33_650M_UR50D (Lin et al. 2023). Subsequently, the deep learning model TropiGAT was built to process the input graph. TropiGAT was designed using an encoder-classifier-like architecture. The encoder module takes the nodes features of B2 and B1, together with their (B2→B1) edges as input and passes it through a GATv2 layer (Brody, Alon, and Yahav 2022). This operation allows the nodes B1 to update their representation by aggregating the embedding representations of the upstream nodes B2. The aggregation is done using a dynamic attention mechanism that attributes to each node B2 an attention coefficient, allowing the associated node B1 to select its most relevant neighbors with respect to the computed loss. As a result, the embedding representations from the set of B2 nodes are effectively compressed into a single 1280-dimensional vector characterizing the associated node B1. The updated features of B1 are then passed into the classifier. The classifier module consists of n linear layers (with n in [2,3]), interspersed by batch normalization, LeakyRelu activation and dropout layers. The raw outputs of the final layer are passed through a sigmoid function, giving an appropriate estimate of the probability of the node B1, given the nodes B2 it is connected to, to be able to infect the KL type of the node A. The architecture was illustrated in **Figure 2A**.

**Figure 2.**
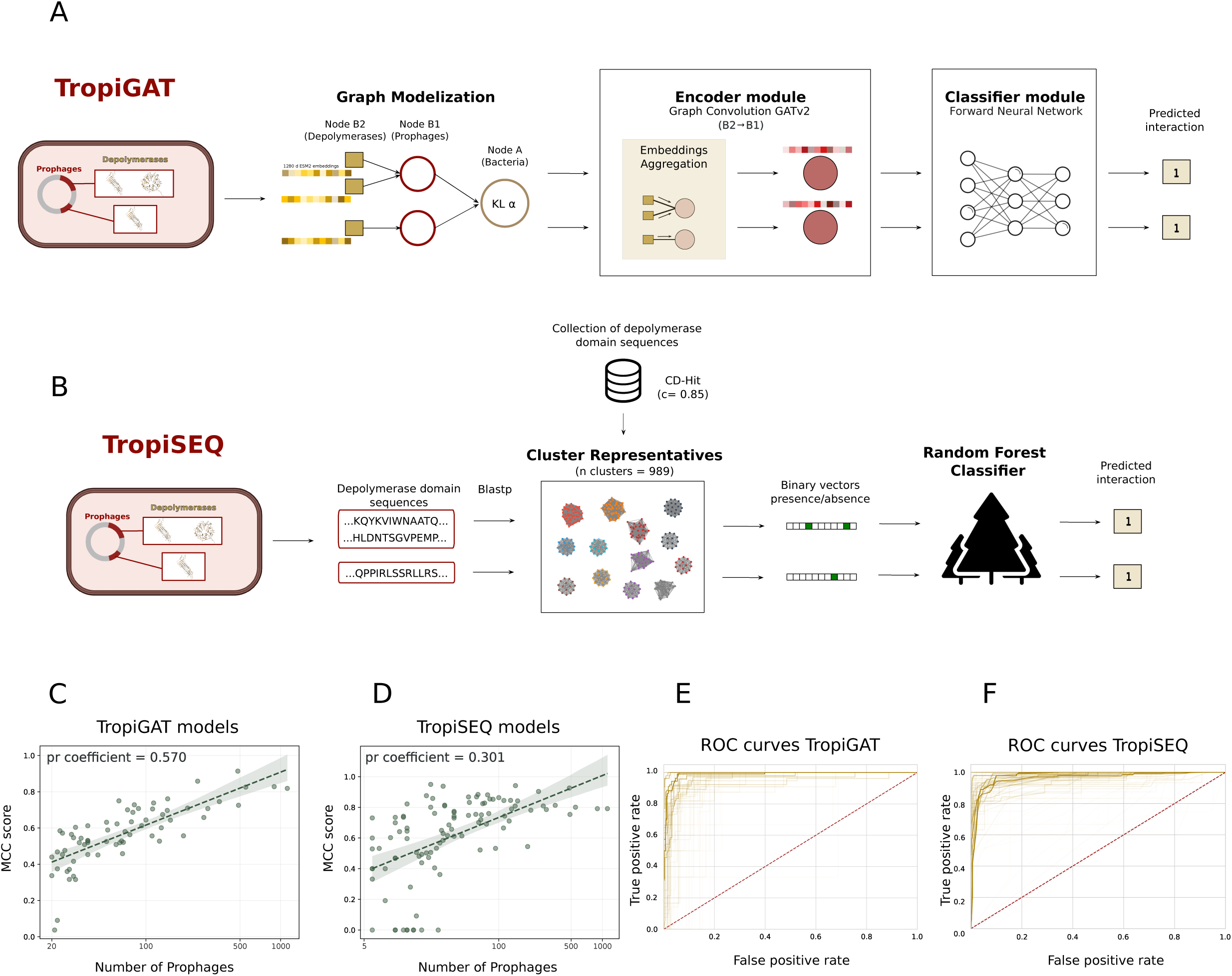
Models employed to predict depolymerase-dependent phage tropism. The elements of the architectures are described using as an illustration the scenario of a lysogen with two prophages, one carrying two de polymerase sequences and the other carrying one 1. **A:** Description of the architecture of TropiGAT. The input is modelized with graphs, where the bacteria, prophage and the encoded depolymerase are represented as nodes and their relationship by edges. The features of the depolymerase are aggregated by the encoder, before being classified. **B:** Description of the architecture of TropiSEQ. The depolymerase domain sequences are retrieved then used to scan the collection of representative depolymerase domain clusters using blastp. The vector representing the presence-absence of each of the depolymerase domain cluster is fed into a Random Forest classifier. **C** : TropiGAT’s and **D**: TropiSEQ’s scatter plots representing the MCC scores on the y-axis along the number of infecting prophages on the x-axis used for training. The green line in bold represents the regression line and the colored area the deviation. ROC AUC curves across the KL types modeled by **E:** TropiGAT and **F:** TropiSEQ. The transparency of the curves was weighted using the number of positive training instances. Pr coefficient = Pearson correlation coefficient.

### Sequence clustering-based approach for modeling prophage-CPS interactions

The second approach for modeling phage-KL type interactions relied on a presence-absence matrix representing the depolymerase domain repertoires within each prophage. To generate this matrix, depolymerase domain sequences identified in the prophages were clustered using CD-HIT, a rapid clustering tool for biological sequences (Li and Godzik 2006). A range of clustering thresholds (0.60, 0.65, 0.70, 0.75, 0.80, 0.85, 0.90, 0.95, 0.975) were assessed, each with a minimal alignment coverage of 80% set by the “-G 0 -aL 0.8” parameters. These thresholds were used to explore the effect of different levels of sequence similarity on the performances of the model. A presence-absence matrix was constructed to represent the depolymerase domain repertoire within each prophage. Each column of the matrix corresponded to a depolymerase domain sequence cluster generated by applying CD-HIT for a specific similarity threshold. The matrix was then used to fit the Random Forest algorithm. The clustering threshold with the best predictive power was determined by computing the weighted Matthew Coefficient Correlation (MCC) for each KL type. The outline of this approach is detailed in **Figure 2B**.

### Training and evaluating the models

An ensemble method was adopted for both modeling approaches based on distinct binary classifiers tailored to each KL type. For each KL type, positive instances were curated by collecting prophages whose infected ancestor possessed the corresponding KL type. To mitigate the risk of data leakage and reduce redundancy, a stringent filtering process was implemented. Prophages were grouped based on their strain, the ancestral bacteria they infected, and their set of depolymerase domain sequences. From each group, only a single representative was retained. This approach addressed the potential issue of nearly identical genomes from bacterial strains sequenced during epidemic outbreaks. The negative instances were randomly selected prophages. To limit the incidence of false negatives, a prophage was considered if none of its depolymerase domain were present in any of the positive instances. The final dataset maintained a ratio of one positive instance to five negative instances.

TropiGAT models were fivefold cross validated using a shuffle split method. At each fold, 70% of the data was used for the training, 20% for the testing and 10% for the final evaluation. TropiGAT instances were generated for KL types with at least 20 prophages, employing a large version with four forward layers in the classifier module for KL types with over 125 positive instances, and a small version with three forward layers otherwise. Early stopping based on MCC stagnation after 60 epochs was implemented, and hyperparameters were optimized using Optuna (Akiba et al. 2019). For TropiSeq, the Random Forest models underwent fivefold cross-validation using a stratified KFold method. Hyperparameters were tuned using a Bayes Search four-fold cross-validation approach through scikit-optimize. The best-performing model, with a c-value of 0.85 resulting in 989 clusters, reached a weighted MCC of 0.776 and was selected for subsequent analysis. The metrics used to evaluate the models were computed along the following formulas: recall (TP / [TP + FN]), accuracy = ([TP + TN] / [TP + TN + FP + FN]), precision (TP / [TP + FP]), F1 score ([precision + recall] / [2 * precision * recall]), area under the ROC curve (AUC), Matthews Correlation Coefficient (MCC; [TN*TP-FN*FP] / sqrt[(TP+FP)*(TP+FN)*(TN+FP)*(TN+FN)]), where TP, FN, TN, and FP refer to true positives, false negatives, true negatives and false positives, respectively. The workflow and the models can be accessed and re-used through the https://github.com/conchaeloko/Klebsiella_TropiSEARCH.

### Prediction on lytic phages

To assess the predictive capability of our models for lytic phages, two datasets were used. The first one consisted of lytic phage depolymerases whose association with KL types were experimentally validated. The second one comprised publicly available infection matrices of *Klebsiella* phages, which reported the phage-bacteria outcome for all the possible interactions (Townsend et al. 2021; Beamud et al. 2023; Ferriol-González et al. 2024). To ensure the identification of all the depolymerase domain sequences in the lytic phages, the three identification methods described earlier were used, supplemented by the information reported by the authors. A phage was considered effective against a given KL type if the authors reported an instance of the phage infecting a bacterium from the corresponding KL type. This criterion accounted for the fact that a depolymerase activity does not necessarily manifest as a visible halo in plaque assays (Sillankorva, Neubauer, and Azeredo 2008; Pires et al. 2016). For TropiGAT models, the architecture was leveraged by making predictions directly from the ESM2 representation of each depolymerase domain sequence. When making predictions with the TropiSEQ models, the amino acid sequence of the depolymerase domains were first scanned against the representatives of the depolymerase domain clusters using blastp. The depolymerase domains were assigned to the cluster of the top hit only if the bitscore was exceeding 75 with a minimal alignment coverage of 80%. Finally, target KL types were inferred from the KL types associated to the corresponding depolymerase domain cluster.

## Results

### A comprehensive profiling of prophage-encoded depolymerases targeting *Klebsiella*

An initial set of 14,601 genomes of *Klebsiella* was downloaded from the NCBI database. Out of these, the KL type was accurately determined for 12,003 genomes, which were retained for downstream analysis. The screening of these bacterial genomes identified 77,802 prophages. To generate a collection of depolymerase domain sequences and the KL type targeted by the prophage encoding them, a bioinformatic pipeline was designed. First, the phylogenetic relationship between the prophages was investigated. A total of 16,077 prophage strains, *i.e* groups of prophages sharing 99% of identity over more than 80% of their genome were identified. For each prophage within a prophage strain, the infected ancestor was deciphered, allowing the inference of the targeted KL type. Out of the 77,802 prophages, 3500 presented an ambiguous target KL type and were discarded, resulting in a collection of 74,302 labeled prophages.

Next, the depolymerase domain sequences within these prophages were screened using three methods. This analysis identified 19,600 depolymerase domain sequences, with 3908 unique sequences (100% coverage and 100% percentage identity). Around 20% (15,230 out of 74,302) of the prophages carried at least one depolymerase. While most prophages harbored a single depolymerase (∼72%), some encoded multiple instances with as many as 12 depolymerases observed in one case. This diversity prompted an average of 1.3 depolymerase sequence per prophage across the database. A final filter was applied to avoid oversampling prophages originating from the same infectious event. This resulted in a final dataset of 8871 prophages, each encoding depolymerase sequences and labeled with one of 128 distinct KL types (**Figure 1C-Supplementary Table 1**). The distribution of prophages was not uniform across the panel of KL types. Indeed, 6 out of 128 KL types were assigned to ∼44% of the prophages in the database: KL107 (1121), KL64 (897), KL47 (551), KL106 (488), KL17 (481) and KL2 (351). About 51% of the prophage database was covered by 61 KL types, each assigned to between 20 and 300 prophages. Lastly, around 5% of the prophage database was covered by 61 KL types, each assigned to fewer than 20 prophages. The investigated folds exhibited varying proportions among the prophages. The most prevalent fold was the right-handed β-helix representing 69% (2722) of the identified sequences, followed by the n-bladed β-propeller at 18% (714). The other folds represented a smaller fraction among which the TIM β/α-barrel with 7.5%, (294), the α/β hydrolase fold with 3% (114), and finally the triple-helix (32) and α/α toroid (29) represented less than 1% (**Figure 1E**).

### Exploring the predictability of prophage tropism

Two approaches were employed to model the complex interplay between bacteria, prophages, and their depolymerase domain. First, TropiGAT relied on a heterograph-based approach where each object is represented by a node and its relationships by edges. The features of the depolymerase domain, represented by embeddings computed with ESM2, were aggregated using an attention mechanism before passing through a deep neural network for binary classification. Second, TropiSEQ relied on a sequence clustering-based approach. The entire set of depolymerase domain sequences retrieved from the prophages was clustered based on sequence homology. Next, representatives of the clusters were used to represent each prophage as categorical features based on their presence-absence in their genome. The resulting vector was finally fed into a Random Forest model for binary classification.

For each KL type, a unique instance of each model was designed and trained on the prophages assigned to the corresponding KL type as positive instances. The performance of the models was evaluated using the MCC. Detailed information about the other metrics can be found in **Supplementary Table 2A-2B**. The performance of TropiGAT was notably influenced by the number of prophages available, as indicated by a Pearson correlation coefficient of r = 0.570 (P < 0.001) between MCC values and the number of prophages. Indeed, the greater the amount of prophage instances associated to a KL type, the better TropiGAT could generalize **(Supplementary Data 1)** and perform on the MCC metric. Specifically, the six most represented KL types had on average a MCC score of 0.810 (standard deviation = 0.067), namely KL2 (0.746), KL17 (0.914), KL47 (0.826), KL64 (0.828), KL106 (0.726), and KL107 (0.819) **(Figure 2C)**. For KL types represented by 120–300 prophages, the average MCC reached 0.672 (standard deviation = 0.093). While some KL types, such as KL102 and KL24, maintained high MCC values 0.859 and 0.707, respectively, others such as KL23 achieved lower MCC (0.528). For the least represented KL types, TropiGAT encountered challenges owing to limited training data, resulting in an average MCC of 0.517 with higher variability (standard deviation = 0.146). This tendency was also observed using ROC curves for assessing model performance **(Figure 2D)**.

In contrast, the performance of TropiSEQ was mildly influenced by the number of representing prophages for a KL type (Pearson r = 0.301, P = 0.002). High and consistent performances were observed for the most represented KL types, with MCC scores ranging from 0.75 to 0.92 **(Figure 2E)**. Despite the decreasing numbers of representative prophages, TropiSEQ maintained the level of performance to an average MCC of 0.796 (standard deviation = 0.104) for the KL types represented by 120-300 prophages. Notably, KL3 (0.942), KL14 (0.846), KL23 (0.711), KL25 (0.809), and KL62 (0.842) exhibited strong performance metrics. TropiSEQ performance slightly decreased for less represented KL types, yet maintaining relatively high scores, with an average MCC of 0.664 (standard deviation = 0.212) for the KL types represented by less than 120 prophages (**Figure 2E)**. The computed ROC curves, validated this trend **(Figure 2F)**.

In conclusion, both approaches exhibited robust predictive capabilities for prophages, with most models displaying MCC scores above 0.5. The number of training instances appeared to be a significant determinant of performance for the models, albeit with a greater impact observed for TropiGAT compared to TropiSEQ.

### Analysis of depolymerase-KL association clusters

TropiGAT utilizes an attention-based graph convolution layer to assign attention weights to each depolymerase domain sequence within a prophage. These weights reflect the model focus on a specific depolymerase domain when predicting the probability of an association between its corresponding prophage and a KL type. By compiling the attention weights attributed to depolymerase domains across all TropiGAT instances, a comprehensive database linking depolymerase domains to specific KL types was generated (**Supplementary Data 2**). Depolymerase domains were considered significantly associated with a KL type if the corresponding attention weight exceeded 0.5 and the predicted probability of prophage infection for that KL type was above 0.8, to ensure high confidence (**Figure 3A**). This approach enabled the mapping of 1760 depolymerase domains to their corresponding KL types, spanning a total of 67 distinct types. Interestingly, 28% (497 out of 1760) of the depolymerase domains exhibited associations with multiple KL types. The most frequent association observed for a single depolymerase domain was KL106-KL107, observed for 11 distinct depolymerases. The following most prevalent combinations were KL64-KL107, KL24-KL112, and KL64-KL106, each involving five depolymerases (**Figure 3B**).

**Figure 3.**
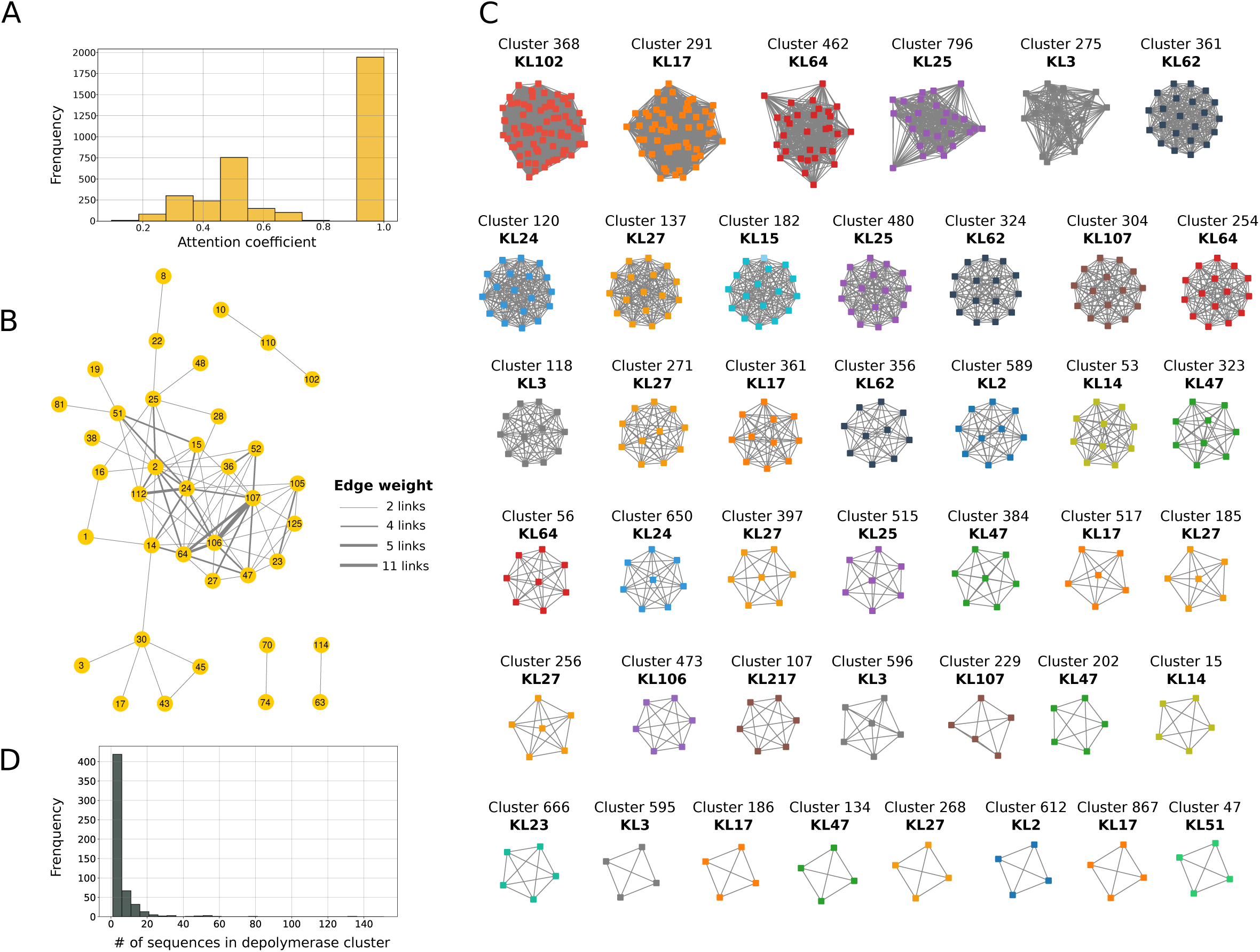
Depolymerase-KL clusters. **A**: Histogram representing the attention coefficient attributed to the depolymerase domain sequences of the training data across the instances of TropiGAT. **B:** Graph representation of the associations between KL types revealed by TropiGAT, with the KL types as nodes and the size of the edge as the prevalence of the association. The sequences that were associated with more than 2 KL types were represented. **C:** Graph representation of depolymerase domain clusters associated with a KL type. Here are represented a subset of the entire set of depolymerase domain sequence cluster for which TropiSeq revealed an association with a probability superior to 0.5. **D:** Distribution of the depolymerase domain cluster sizes that presented an association with at least one KL type.

To assess the association between depolymerase domain clusters and KL types using TropiSEQ, predictions were performed on vectors representing individual depolymerase domain clusters. This enabled the determination of the probability of the interaction between each depolymerase domain cluster with the KL types. A probability score threshold of 0.5 was applied, resulting in the identification of 550 (out of 989) depolymerase domain clusters associated with 96 distinct KL types (Figure 3C). Among these, 110 (20%) clusters displayed associations with at least two KL types (Supplementary Data 2). The most common associations were KL106-KL107 with seven depolymerase domain clusters, followed by KL47-KL64 with five clusters in common. Most associated clusters were constituted of few depolymerase domain sequences, with the most prevalent size being singletons with 180 clusters (out of 550), followed by clusters with two depolymerase domain sequences (115 out of 550; **Figure 3D**). The largest clusters contained 156 and 153 sequences.

The KL type labels assigned to depolymerase domains by TropiGAT were compared with those assigned to their clusters by TropiSEQ. Out of 1896 depolymerase domain-KL type combinations revealed by TropiGAT, 1645 (86.8%) were congruent with the assignments made by TropiSEQ.

### Applicability of the models for predicting the KL tropism of lytic phage depolymerases

To assess the applicability of the models to *Klebsiella* lytic phages, a dataset of experimentally validated depolymerases was compiled. It comprised 25 pairs of right-handed β-helices with their target KL type (**Supplementary Table 3**). To evaluate the performances of TropiGAT and TropiSEQ, all their instances were used to make predictions. The generated probabilities of interaction between depolymerases and each KL types were then sorted. Two key factors determined successful prediction: (1) whether the correct KL type received a probability exceeding 0.5 and (2) its relative value of their associated probabilities with other KL types.

Among the 25 tested depolymerase-KL type pairs, both TropiGAT and TropiSEQ correctly identified 11 (**Table 1**). At least one model successfully predicted 15 pairs (60%). The majority of successful predictions (9/15) overlapped between the two methods. For example, KL47 (recall: 4/4 for both TropiGAT and TropiSEQ) and KL64 (recall: 3/3 for TropiGAT, 2/3 for TropiSEQ) were consistently predicted within the top five predictions. Interestingly, TropiGAT identified a potential cross-reaction between KL2 and KL13 by predicting positive interactions for both KL types in *Klebsiella* phage vB_KpnP_KpV74 (Solovieva et al. 2018; Pilar Domingo-Calap et al. 2020). Analysis of missed predictions (n=10) indicated a lack of significant sequence identity between the tested depolymerases and the training data (n=5), or an association with a depolymerase cluster that was not associated with any KL type (n=4). Neither KL1 (recall: 0/4) nor KL102 (0/1) were correctly predicted, supporting previous research that identified a lack of similarity between their depolymerases and those found in prophages (Beamud et al. 2023).

**Table 1.** Performances of the predictive models on the set of experimentally validated depolymerases. “-” : not predicted.

| KL types | TropiGAT |  | TropiSEQ |  |
| --- | --- | --- | --- | --- |
|  | recall | ranks | recall | ranks |
| KL47 | 4/4 | 1;1;1;5 | 3/4 | 1;1;-;1 |
| KL1 | 0/4 | - | 0/4 | - |
| KL64 | 3/3 | 3;4;5 | 3/3 | 1;1;1 |
| KL2 | 1/3 | 3;-;- | 0/3 | - |
| KL21 | 0/2 | - | 1/2 | 2;- |
| KL13 | 1/1 | 9 | 1/1 | 1 |
| KL30 | 1/1 | 4 | 1/1 | 1 |
| KL3 | 1/1 | 4 | 0/1 | - |
| KL35 | 0/1 | - | 1/1 | 1 |
| KL63 | 0/1 | - | 1/1 | 1 |
| KL11 | 0/1 | - | 0/1 | - |
| KL25 | 0/1 | - | 0/1 | - |
| KL102 | 0/1 | - | 0/1 | - |
| KL69 | 0/1 | - | 0/1 | - |

To gain more insights about the performances of the models in lytic phages, a comprehensive dataset was generated from three infection matrices publicly available. In the matrix by Ferriol-Gonzales et al, 2024, interactions between 71 phages and 77 bacteria covering as many different KL types were presented. The context of that study was the isolation of phages capable of infecting the full diversity of the 77 serotypes in *Klebsiella*. In the second matrix, Beamud et al, 2023 described the outcome of the interaction between 46 isolated phages and 138 bacterial strains covering 59 different KL types. There, the bacterial strains were sequenced in a clinical context. Finally, Townsend et al, 2021 presented 30 phages isolated tested against 24 bacterial strains covering 18 different KL types. Overall, 89 different KL types were covered in the matrix. Each phage within these matrices was re-annotated for the depolymerase domain sequence using the pipeline previously described. The re-annotation resulted in the identification of 249 depolymerase domain sequences in 126 phages targeting bacteria for which the KL type was predicted with a good level of confidence. The depolymerase domain folds consisted of 139 right-handed β-helix, 75 n-bladed β-propeller and 34 triple-helix and 1 α/α toroid **(Supplementary Table 4)**. For each depolymerase domain, predictions were made using all the instances of TropiGAT and TropiSEQ.

Out of 328 reported phage-KL type interactions, TropiGAT could predict 115 (∼35%), TropiSEQ 49 (∼15%) and 133 (∼40%) when both models combined (**Figure 4A**; **Supplementary Table 5**). As expected, the proportion of the phage-KL type combination that correctly predicted was lower than for the dataset of experimentally validated depolymerase (∼60%). This difference could be the consequence of the phage-bacteria interactions that are not mediated by the depolymerase, consequently not detected by the models.

**Figure 4.**
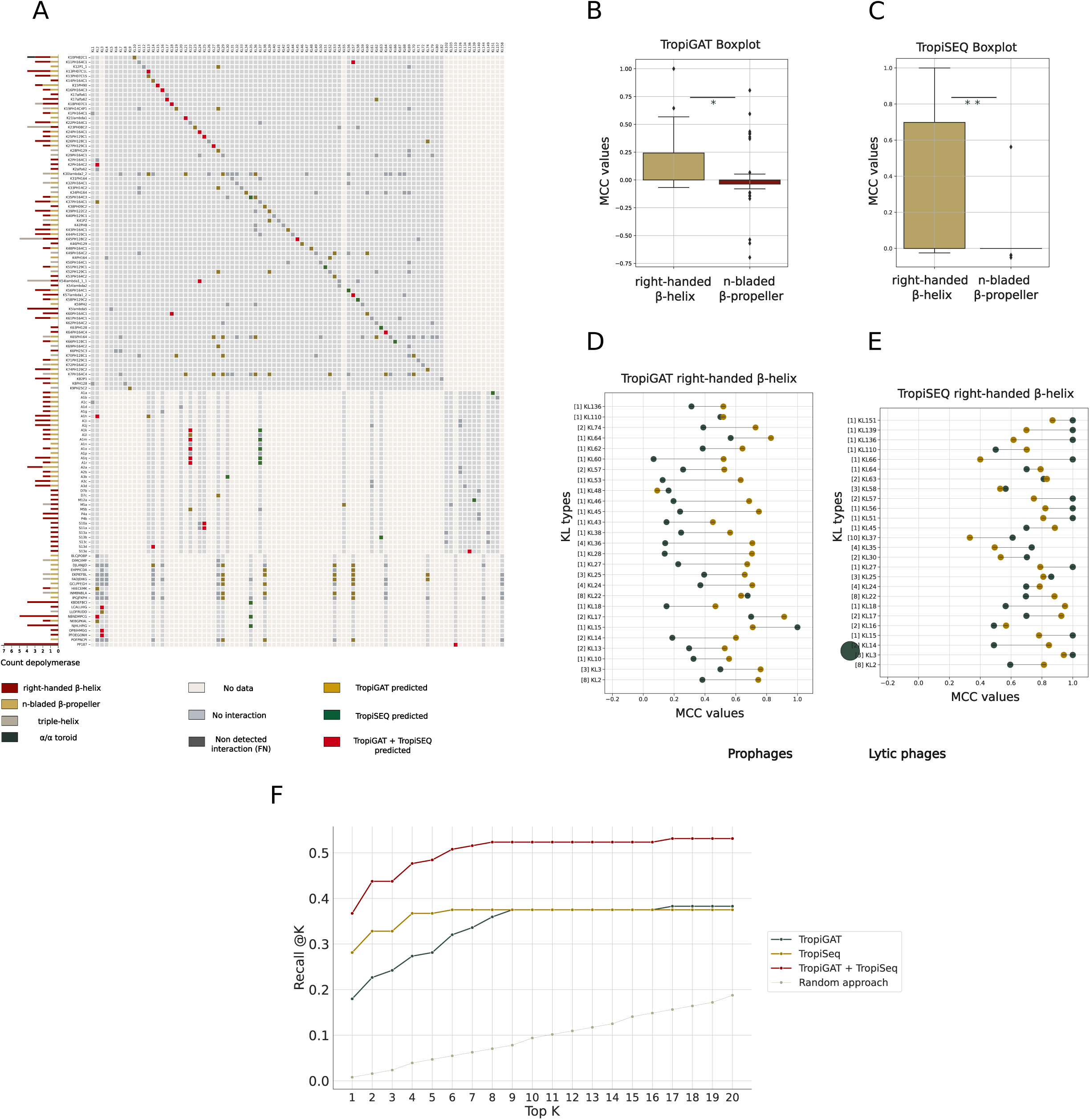
Model performance on lytic phages. **A**: Compilation of the three experimental infection matrices used for evaluating the models. In the x-axis are represented KL types and in the y-axis their corresponding phages. In white: combination not tested; gray: no observable infection; dark gray: infection not predicted by neither of the predictors; green: infection predicted by TropiSEQ; golden: infection predicted by TropiGAT; red: infection predicted by both models. Box plot of the MCC values reported for the KL types when only considering the right-handed β-helix and n-bladed β-propeller when predicting with **B**: TropiGAT and **C:** TropiSEQ. * : <0.05, **: <0.005. Dumbbell plot of the MCC value for the different KL types calculated for the lytic phages for the **D**: TropiGAT and **E**: TropiSEQ models. **F**: Recall @K for TropiGAT (green), TropiSEQ (gold), the combination of TropiGAT and TropiSEQ (red) and the random approach (gray).

For most of the KL types with large amount of instances in the training data, the performances of the models transferred successfully from the prophages to the lytic phages when considering the right-handed β-helix. For example, TropiGAT and TropiSEQ presented MCC scores of 0.828 and 0.791 in prophages and 0.567 and 0.701 in lytic phages respectively for KL64 (**Figure 4D**, **Figure 4E)**. Overall, 27 KL types presented an MCC score of at least 0.4 for one of the two approaches. Moreover, both TropiGAT and TropiSEQ could specifically predict some KL types. Interestingly, for TropiSEQ those KL types tended to present very few instances in the training data, like KL37 (6), KL56 (13), KL58 (6) or KL66 (8) suggesting that despite low training representation, TropiSEQ can present good predictive abilities applicable to lytic phages.

Overall, the results indicate that the models built on lysogenic phages have potential for predicting KL tropism in lytic phages based on the right-handed β-helix, particularly for KL types well-represented by depolymerases in the training data (e.g., KL47 and KL64). The degree of sequence homology between the tested depolymerase and those encoded by prophages the training data appears to be a critical factor for successful prediction.

### The depolymerase fold impact the prediction of capsular specificity

The right-handed β-helix has been the most extensively characterized depolymerase fold, while the characteristics and properties of other folds are less understood. The previous section has reinforced the idea that KL type tropism can be predicted for the right-handed β-helix, but the predictability of other folds remains unclear. In order to explore these factors, the predictability of the different depolymerase folds was investigated. The results indicated that the performances of the model were not equal across the folds, as the right-handed β-helix harnessed 49 and 48 correct depolymerase domain-KL type combinations, and the n-bladed β-propeller 48 and 1, for TropiGAT and TropiSEQ respectively. To investigate this further, the MCC scores of the predictions built on prophages were computed based on each fold and compared. For both TropiGAT and TropiSEQ, clear discrepancies were observed for the MCC scores calculated based on the right-handed β-helix versus the n-bladed β-propeller. For TropiGAT, the difference was notable with an average MCC score of 0.135 for the right-handed β-helix and 0.001 for the n-bladed β-propeller (P < 1.10e-3; **Figure 4B)**. For TropiSEQ, the differences were even more pronounced with MCC scores of 0.333 and 0.007 for predictions based on the right-handed β-helix and the n-bladed β-propeller, respectively (P < 1.10e-4; **Figure 4C)**. These results indicate that the models are more efficient at making predictions on the right-handed β-helix compared to the n-bladed β-propeller.

Despite the overall lower predictability of the n-bladed β-propeller, the models showed accurate predictions for certain KL types, suggesting some specificity in the predictive capability. This was particularly the case for KL21 that presented MCC scores of 0.370 and 0.562 for TropiGAT and TropiSEQ respectively. TropiGAT also presented good scores for KL16 (0.435), KL27 (0.412), KL38 (0.387), KL52 (0.806) and KL57 (0.594) using the n-bladed β-propeller (**Supplementary Figure 1**). These findings suggest that KL type tropism predictability for n-bladed β-propeller folds may be possible for specific KL types. Interestingly, the basis of this predictability seems to transcend mere protein sequence identity, potentially hinging on specific residues and structural features undetected by sequence-based methods like TropiSEQ. This observation calls for further investigation to clarify the underlying mechanisms and refine predictive capabilities across various depolymerase folds.

### Extending the application of the models: designing a ranking system

Predictive models for phage-bacteria interactions are valuable tools, particularly for characterizing phage host range. To evaluate the models as recommenders for phage-bacteria probing, a ranking system was designed. Here, the focus was put on the right-handed β-helix due to its predictive power discussed previously. For a depolymerase domain, the system takes as an input the probabilities for the KL types being a target and outputs a ranked list. Performance in identifying the correct KL type within the top K predictions (where K is an integer) was evaluated for depolymerase domains from lytic phages. For this, the number of correctly predicted target KL types in the top K predictions for each depolymerase domain divided by the total number of observed phage-KL type interactions. Alongside TropiGAT and TropiSEQ, the performance of a random approach was designed using a Monte Carlo simulation (Harrison, Granja, and Leroy 2010).

TropiSEQ demonstrated great abilities to identify the correct KL type within the top predictions for low values of K. At K=1 (*i.e.* considering only the top prediction), TropiSEQ correctly predicted interactions for 36 phage-KL types pairs (∼28% of total phage-KL type interactions), compared to 23 (∼18% of total phage-KL type interactions) for TropiGAT and only two using a purely random approach **(Figure 4F)**. TropiSEQ performances peaked at K=6, where all the 48 correct predictions were made, while the random approach reached seven correct predictions. The random approach becomes equivalent to TropiSEQ at K=39. In contrast, the recall @K steadily increased for TropiGAT until reaching a plateau at K=9 **(Figure 4F)**. Hence, TropiSEQ demonstrated great abilities for identifying the correct KL type within the very top predictions, while TropiGAT can provide more information when considering a broader range of potential KL types. This highlights the potential benefit of combining the outputs of both models when aiming to comprehensively characterize the spectrum of KL types a phage might interact with.

The graph-based approach of TropiGAT leverages embedding representations, capturing not only sequence similarity but also folding patterns of depolymerase domains. This empowers TropiGAT to make precise predictions on lytic phage depolymerase domain sequences, even when they lack significant homology with any training data. Notably, TropiGAT successfully assigned target KL types to depolymerase domains that eluded TropiSEQ (**Table 2**). For example, it correctly predicted interactions between KL10 and the phage vB_Kpn_K10PH82C1 (right-handed β-helix on CDS 50, probability = 0.956, rank 2) and between KL53 and the phage EKPIEFBL (triple-helix on CDS 113, probability = 0.981, rank 2) despite a lack of significant sequence homology in their respective depolymerase domains with the sequences of the training data. TropiGAT ability extended to domains where TropiSEQ offered no predictions, such as those of phage vB_Kpn_K60PH164C1 interacting with KL60 (right-handed β-helix on CDS 96, probability = 0.996, rank 1) and KL18 (right-handed β-helix on CDS 94, probability = 0.999, rank 1; **Table 3**). These findings suggest TropiGAT leverages patterns beyond sequence similarity. By capturing folding patterns, the embeddings enable the model to learn and make correct inferences at a higher structural level. Overall, these results demonstrate that the ranking system is a valuable tool when characterizing phage depolymerase tropism, allowing the identification of candidates KL types for depolymerase sequences.

**Table 2.** Correct predictions made with TropiGAT, on depolymerase domain sequences with no close sequence homology with the depolymerase domain of the training database.

| Phage | Protein | Folds | TropiGAT correct calls | Ranks |
| --- | --- | --- | --- | --- |
| EKPIEFBL | cds 113 | triple-helix | KL53 | 8 |
| FADJDIKG | cds 19 | triple-helix | KL53 | 14 |
| INMBNBLA | cds 202 | triple-helix | KL53 | 8 |
| K10PH82C1 | cds 50 | right-handed β-helix | KL10 | 2 |
| K19PH14C4P1 | cds 48 | triple-helix | KL28, KL19 | 11, 13 |
| K2064PH2 | cds 25 | triple-helix | KL28, KL74 | 12, 10 |
| K28PH129 | cds 24 | triple-helix | KL28 | 14 |
| K33PH14C2 | cds 25 | triple-helix | KL28 | 10 |
| K52PH129C1 | cds 25 | triple-helix | KL28, KL39 | 11, 9 |
| LLOFRUDD | cds 39 | triple-helix | KL3 | 12 |
| NBNDMPCG | cds 165 | right-handed β-helix | KL2 | 14 |

**Table 3.** Correct predictions made with TropiGAT, on depolymerase domain sequences for which TropiSEQ could not make any inferences.

| Phage | Protein | Folds | TropiGAT correct calls | Ranks |
| --- | --- | --- | --- | --- |
| K17alfa62 | cds 66 | right-handed β-helix | KL62 | 1 |
| K26PH128C1 | cds 49 | right-handed β-helix | KL74 | 1 |
| K37PH164C1 | cds 48 | right-handed β-helix | KL2 | 1 |
| K38PH09C2 | cds 24 | right-handed β-helix | KL38 | 1 |
| K45PH128C2 | cds 237 | right-handed β-helix | KL45 | 9 |
| K53PH164C2 | cds 24 | right-handed β-helix | KL53 | 2 |
| K60PH164C1 | cds 96 | right-handed β-helix | KL60 | 1 |
| K60PH164C1 | cds 94 | right-handed β-helix | KL18, KL60 | 1,2 |
| K74PH129C2 | cds 51 | right-handed β-helix | KL74 | 1 |
| PP187 | cds 230 | right-handed β-helix | KL110 | 1 |
| K13PH07C1S | cds 11 | right-handed β-helix | KL13 | 6 |

## Discussion

This study proposes an innovative approach to the important, yet difficult task of modeling the outcome of phage-bacteria interactions at the sub-species level in *Klebsiella* spp. By leveraging the data contained in prophages, models of phage capsular tropism based on their respective depolymerase domains were generated. These models built using prophage data demonstrated predictive abilities that could be transferred to lytic phages. Additionally, a comprehensive database of depolymerase domain sequences associated with their target KL types was compiled.

Predictive performances of the models were unequal across the KL types. A clear positive correlation was observed between the number of prophage instances labeled with a KL type and the corresponding performance of the model. For some KL types, models could not be generated due to insufficient amount of training data. Nevertheless, this does not diminish the model relevance, especially when considering the clinical context as high-risk clones of *Klebsiella* spp. are known to be associated with specific KL types (Wyres, Lam, and Holt 2020). By specifically collecting *Klebsiella* strains with relevant KL types from patients, researchers can enrich collections with these prophages, ultimately increasing training datasets for those KL types. Indeed, the most represented KL types in this study are the ones reported in the literature as clinically relevant, namely KL2, KL17, KL47, KL64, KL106 and KL107 (Cai et al. 2022; J. Wang, Feng, and Zong 2023). Additionally, other clinically relevant bacteria such as *Escherichia coli* or *Acinetobacter baumannii* express a capsule encoded by the Wzy system and are targeted by phages encoding depolymerase domains (Abdelkader et al. 2022; Blasco et al. 2022). Thus, the approach used here can be generalizable to these species as well as representing a potential pool of new informative patterns for the models (Pas et al. 2023; Yiyan Yang et al. 2024).

Evaluation of the models on the lytic phages revealed varying levels of predictive power across the different depolymerase domain folds, prompting questions on the origins of these discrepancies. The right-handed β-helix is the most described polysaccharides degrading depolymerase in phages due to its ubiquity and stable trimer structure (Shneider et al. 2013; Buth et al. 2015), typically targeting one or few different KL types (Y.-J. Pan et al. 2017). In contrast, while ongoing research explores the properties of the n-bladed β-propeller in bacteria and eukaryotes (Saccuzzo et al. 2024; C. Chen et al. 2024), its characteristics in phages remain unclear. Several factors could account for the weak predictive power of this fold. Firstly, the scarcity of training data, as the detectable n-bladed β-propeller is less abundant than the right-handed β-helix in phages (Concha-Eloko et al. 2024). Secondly, recent studies demonstrated vast structural diversity within this fold. The β-propeller structure results from the duplication of four antiparallel β-strands, which can then act as a monomer or dimer, leading to variation in the number of blades (Pereira and Lupas 2022). This diversity could complicate the identification of pattern. Finally, the n-bladed β-propeller can accommodate a wide range of substrates depending on environmental conditions (Pandey et al. 2021), exerting diverse functions that can go beyond catalytic activity (C. K.-M. Chen, Chan, and Wang 2011). These features align with the findings of this study, as phages encoding an n-bladed β-propeller tended to target more KL types than those lacking it, with an average of 3.1 versus 1.9 targets respectively, suggesting a broader infectivity range. Further research into the precise activity of this fold in phages is needed to gain deeper insights.

Focusing exclusively on the depolymerase domain sequence in modeling phage-bacteria interactions has limitations. The recognition process can be mediated by the tail fiber and occur independently of the depolymerase (Concha-Eloko et al. 2023), in conjunction with it (Dunstan et al. 2021) or alongside it, particularly in phages exhibiting depolymerase activity with specific KL types but not others. The complexity of the phage recognition system could lead to an overestimation of false negatives, to the extent that successful infections may not always involve depolymerase activity, despite the phage encoding one. Moreover, the ability of a phage to recognize its host does not necessarily lead to successful infection due to post-entry defense mechanisms. This scenario could lead to potential false positives in cases where a phage can exert depolymerase activity but fails to cause a successful infection.

To address these limitations, models could be expanded by integrating other factors involved in the recognition of the host. This could entail integrating tail fibers and the secondary bacterial receptors. Recent genomic foundation models capable of generating embedding representations for DNA sequence were developed with hundreds of thousands of kilobases (Nguyen et al. 2024). These tools could be leveraged to encode the whole K-loci and allow the incorporation of more informative data about the capsule and its modifications. Moreover, the model could be extended by integrating data on downstream processes such as phage replication, host defense systems (e.g CRISPR), and agents involved in bacterial lysis (e.g holin, endolysin) (Lood et al. 2022). This would allow TropiGAT to predict more comprehensively the success of phage infection, making it a valuable tool for various applications, particularly in clinical settings.

The rationale of the study rested on the assumption of equivalence or continuity between the depolymerase domain sequences found in prophages and lytic phages. The validity of this hypothesis was evidenced by the ability of the model to make accurate predictions for lytic phages using training data from prophage depolymerase domains. Yet, a decline in performance was observed when transferring the domain of prediction from prophages to lytic phages, with variations across KL types. These variations could be attributed to differences in the genomic environments of the lytic and prophage depolymerase domains. Lytic phage depolymerases are transferred along the phage during active replication, whereas prophage depolymerases reside in a dormant state within the host genome. These distinct contexts may result in different selection pressures on their sequences, leading to variations between the depolymerases of these two phage types. To bridge these differences, fine-tuning could be employed (Hosna et al. 2022), which allows a model to adapt to a new dataset after the initial training.

In summary, this study suggests new approaches to modeling phage-bacteria interactions by combining advanced machine learning models and architectures with an evolutionary perspective. The significant role and versatility of the depolymerase sequence enables the scope of this work to go beyond the therapeutic applications, offering potential utility in industrial contexts as well.

## Acknowledgments

This work was financially supported by Advanced Grant 101019724 - EVADER from the European Research Council (ERC) to R.S; fellowhsip GRISOLIAP/2020/158 from the Conselleria d’Innovació, Universitats, Ciència i Societat Digital (Generalitat Valenciana) to R.S. and R.C.-E; P.D.-C was funded by ESCMID Research Grant 20200063, project PID2020-112835RA-I00 funded by MCIN/AEI/10.13039/501100011033, and project SEJIGENT/2021/014 funded by Conselleria d’Innovació, Universitats, Ciència i Societat Digital (Generalitat Valenciana); B.B was funded by a PhD fellowship from Spanish MCIU FPU16/02139.

## Author contributions

Conceptualization & Methodology: R.C.-E., B.B., P.D.-C., R.S.;

Data Curation, Software, Formal Analysis, Validation & Original Draft preparation: R.C.-E.;

Manuscript Review & Editing: R.C.-E., B.B., P.D.-C., R.S.;

Supervision: B.B., P.D.-C., R.S..

All authors read and approved the final manuscript. The authors declare no competing interests

## Data availability

The csv file containing the data used for the training and the evaluation of the models can be accessed through the Zenodo accession: 10.5281/zenodo.1279778.

## Supplementary Material

**Supplementary Figure 1.** Dumbbell plot of the models’ performances on prophages and lytic phages. **A**: TropiGAT’s and **B**: TropiSEQ’s performances on the right-handed β-helix. **C**: TropiGAT’s performances on the n-bladed β-propeller.

**Supplementary Algorithm 1.** Algorithm used to assess the prophages’ infected ancestor to infer the KL type at the movement of infection.

**Supplementary Data 1.** Plots of the train and test loss across the epochs during the training process of TropiGAT for each KL type.

**Supplementary Table 1.** Compilation of the training data, comprising the depolymerase sequences, the encoding prophage, the KL type of the infected ancestor and related information.

**Supplementary Table 2. A.**TropiGAT’s and **B.**TropiSEQ performances on the evaluation dataset of the prophage data.

**Supplementary Table 3.** Detailed information of the experimentally validated depolymerases.

**Supplementary Table 4.** Detailed information of the matrices, depolymerase domain identified and their fold.

**Supplementary Table 5.** Predictions of the models TropiGAT and TropiSEQ on the lytic phages.

