## Supplementary material for "Unlocking data in *Klebsiella* lysogens to predict capsular type-specificity of phage depolymerases": Supplementary_Algorithm_1.pdf

**Data:**

- *Targets*: list of bacteria indices carrying the instances of a prophage strain.
- *Lysogen*: an element in the *Targets* list.
- *LCA*: index of the last common ancestor of the *Targets*.
- *Ancestors*: ordered list of the ancestral nodes going from *Lysogen* to *LCA*.

**Result:** Identify the KL type of the *ancestral bacteria* infected by the prophage present in *Lysogen*.

```
for ancestral bacteria in Ancestors do
    clades: list of clades connected to ancestral bacteria;
    n = 0;
    for clade in clades do
        if clade contains at least one leaf present in Targets then
            n = n + 1
        else
            Continue
        end
    end
    if n > 1 then
        Continue
    else
        infected ancestor = ancestral bacteria;
        return the KL type of the infected ancestor
    end
end
```

**Algorithm 1:** Recovering the KL type of the *infected ancestor*
